# Protocol for scalable generation of uniform 3D porcine oviduct epithelial cell spheroids

**DOI:** 10.64898/2026.06.11.731731

**Authors:** Leonardo M. Molina, Hao-Chun Fan, Adrienne M. Antonson, David J. Miller

## Abstract

While previous *in vitro* porcine (*Sus scrofa*) oviduct spheroid models often rely on manual selection of the formed aggregates, here, we establish a scalable and reproducible system for generating oviduct spheroids. We describe steps for isolating primary secretory and ciliated oviduct epithelial cells and their assembling them into spheroids by forced aggregation using the AggreWell platform. This enables the mass production of uniformly sized spheroids ready for use in a range of downstream applications, such as sperm-oviduct co-incubation experiments.

For complete details on the use and execution of this protocol, please refer to Molina et al.^1^

**Graphical abstract:** 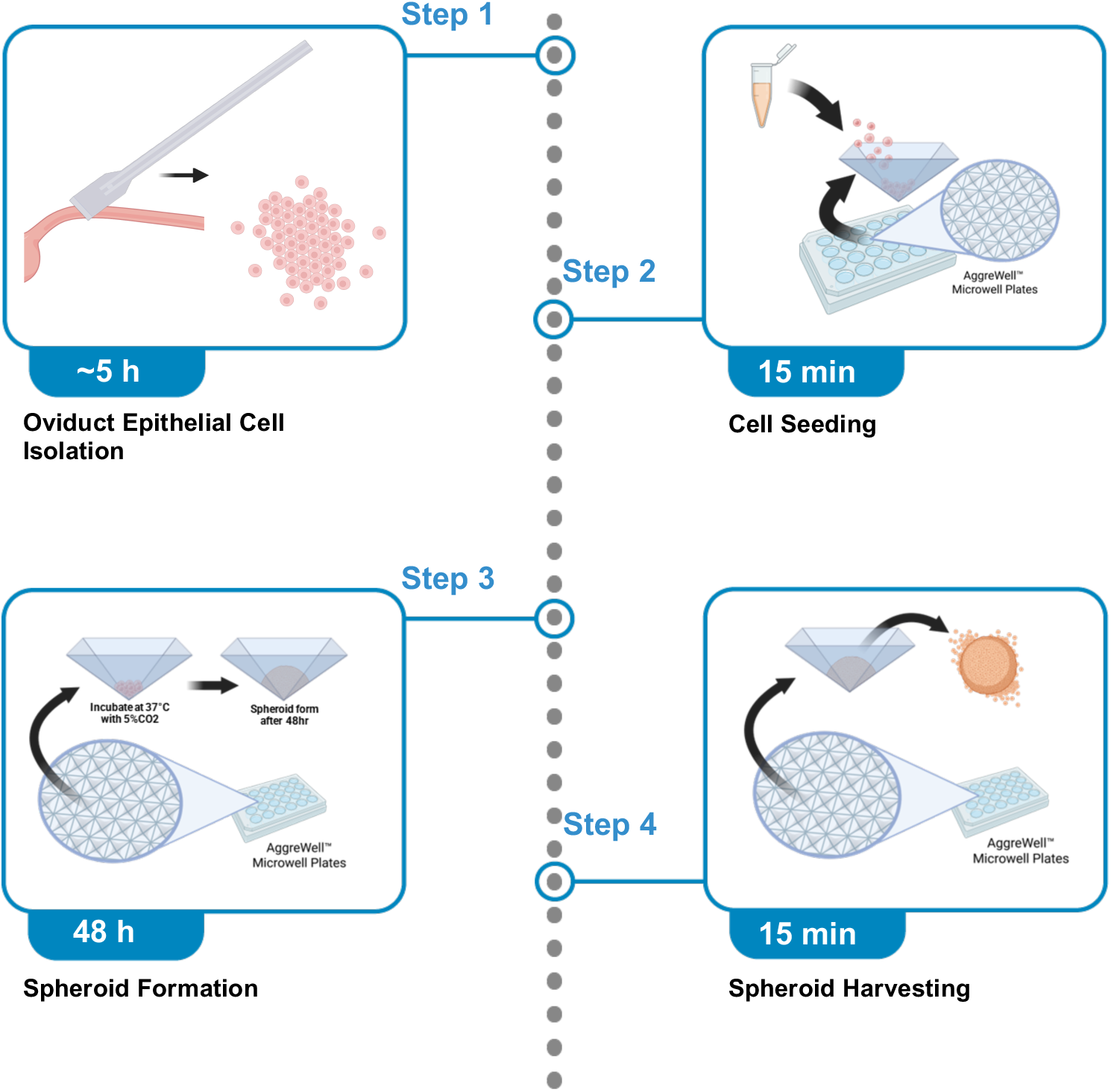

**Before you begin:** This protocol describes the isolation of primary porcine oviduct epithelial cells and their use in generating a three-dimensional (3D) spheroid model *in vitro*. The protocol is specifically designed to enrich and preserve ciliated oviduct epithelial cells, which are critical for physiologically relevant sperm-oviduct interaction studies yet are frequently underrepresented in conventional *in vitro* oviduct models. To minimize the de-differentiation of ciliated cells during culture, a specialized Incubation Medium is employed to maintain epithelial cell viability and ciliary function. The relatively brief spheroid formation period (48 h) further supports preservation of ciliary structure and activity prior to downstream functional applications.

This protocol consists of four major steps. Steps 1 and 2 cover oviductal epithelial cell isolation through mechanical and enzymatic digestion, step 3 describes the spheroid formation and culture conditions, and step 4 applies the spheroid model to sperm co-incubations or immunofluorescent staining. Although optimized for the porcine isthmus, this protocol may also be suitable for the cells from the ampulla and adaptable to other large animal species.

## Innovation

Studying the oviduct *in vitro* has historically presented significant challenges. Conventional two-dimensional (2D) cell culture systems promote rapid de-differentiation of ciliated epithelial cells and loss of cell polarity, limiting the physiological relevance of these models^2,3^. Because sperm-oviduct interactions are partially dependent on ciliary function, the progressive loss of cilia in 2D cell cultures complicates the interpretation of functional studies.

To address these limitations, several 3D oviduct models have been developed, including spheroid cultures, organoids, and air-liquid interface (ALI) systems. However, each approach carries practical drawbacks that can hinder their implementation. ALI cultures require specialized equipment for monitoring, extended culture periods, and depend on cellular re-differentiation to restore ciliation^4^. Oviduct organoids are typically grown embedded within an extracellular matrix, resulting in a single-cell-layer structure with apical polarity directed toward a central lumen^5–8^. This configuration limits direct luminal access and complicates assays that depend on apical surface exposure, including sperm binding and bacterial or viral infection models. Existing spheroid protocols for livestock species offer a more accessible alternative^9,10^; however, they commonly rely on manual selection of aggregates based on size and morphology, a process that is labor-intensive and limits scalability.

Here, we present a standardized protocol for generating oviduct epithelial cell spheroids from primary porcine cells by forced aggregation in microwells, implemented here using AggreWell^TM^ plates. In this approach, dissociated cells are confined within an array of uniform microwells and centrifuged to drive their aggregation into discrete, evenly sized spheroids, eliminating the reliance on spontaneous aggregation and manual selection. By combining established isolation methods for ciliated and secretory epithelial cells^4^ with controlled microwell-based aggregation, this method enables the reproducible production of hundreds to thousands of spheroids with uniform size and cellular composition, facilitating functional assays including sperm co-incubations, infection models, and hormone response studies.

## Institutional permissions

Porcine reproductive tracts were obtained immediately postmortem from animals processed for food production at a local commercial abattoir. No live animals were used in the establishment of this protocol; therefore, approval by the University of Illinois Urbana-Champaign Institutional Animal Care and Use Committee was not required. If the use of primary cells derived from live animal sources is intended, approval from the respective local and institutional animal ethics committees may be required to ensure compliance.

## Oviduct Epithelial Cell Isolation

### Timing: 1 h from slaughter

1. Transport porcine female reproductive tracts to the lab at room temperature within 1 h from slaughter.

**Note:** For this protocol, tracts are transported at room temperature (∼22°C–25°C), which consistently yielded high-quality oviduct epithelial cells when processed within the 1 h window. While alternative transport methods (such as immersion in protective media or preservation on ice) were not evaluated for this specific spheroid protocol, they may be explored by the user if extended transit times are unavoidable.

**CRITICAL:** The following procedures have been optimized for primary cells derived from porcine pre-pubertal gilts. Although potentially applicable to post-pubertal gilts, sows, and other mammalian species, the use of sexually mature tissues may compromise the yield and quality of cells.

Reproductive tracts selected for this protocol should exhibit a prepubertal phenotype: small in size, pale pink to light red in coloration, minimally vascularized (avoiding congested, deep purple, or heavily bruised tissues), only small follicles, and no corpora lutea (**Figure 1A**).

**Figure 1.**
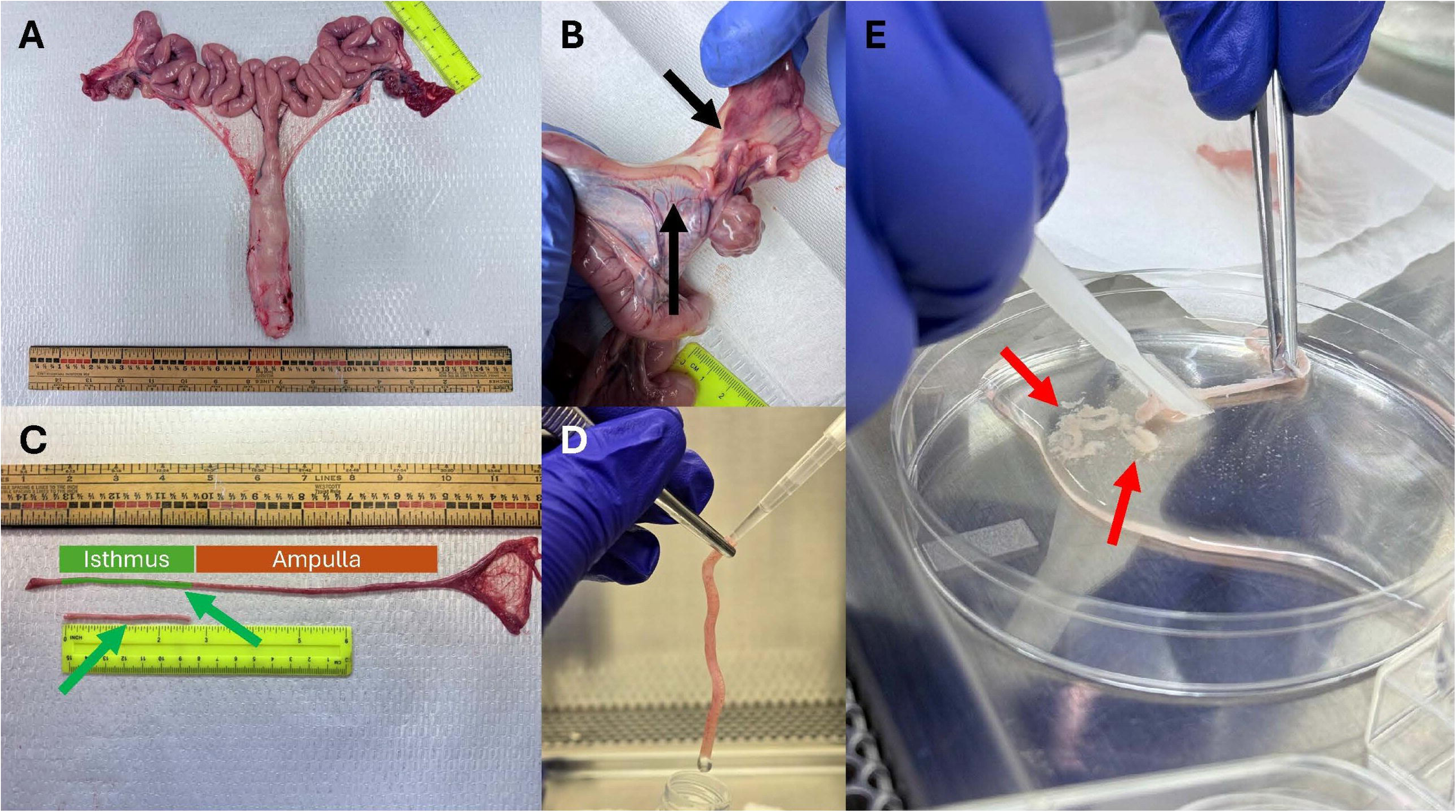
Dissection of the reproductive tract and isolation of oviduct epithelial cells. Representative images showing the isolation of porcine oviduct epithelial cells: (A) orientation of the reproductive tract prior to oviduct dissection, (B) removal of surrounding connective tissue (black arrows), (C) isolation of the isthmus region (highlighted green/green arrows) of the oviduct, (D) flushing of the oviduct lumen prior to collagenase treatment, and (E) extrusion of oviduct epithelial cells (red arrows) from the lumen following collagenase treatment.

## Key resources table

| REAGENT or RESOURCE | SOURCE | IDENTIFIER |
| --- | --- | --- |
| **Antibodies** |  |  |
| Mouse monoclonal anti-acetylated α-tubulin (clone 6-11B-1) | Sigma-Aldrich | Cat# MABT868;<br>RRID:AB_2819178 |
| Mouse monoclonal anti-cytokeratin pan (mixture) | Sigma-Aldrich | Cat# C2562;<br>RRID:AB_476839 |
| Rabbit monoclonal anti-vimentin (D21H3) | Cell Signaling Technology | Cat# 5741;<br>RRID:AB_10695459 |
| Rabbit polyclonal anti-E cadherin | Biorbyt | Cat#orb213706 |
| Rabbit polyclonal anti-PAX 8 | Proteintech | Cat# 10336-1-AP;<br>RRID:AB_2236705 |
| Rabbit polyclonal anti-ESR1 | ABclonal | Cat# A0296;<br>RRID:AB_2757108 |
| Rabbit polyclonal anti-PGR | ABclonal | Cat# A0321;<br>RRID:AB_2757125 |
| Mouse IgG isotype control | Thermo Fisher Scientific | Cat# 08-6599;<br>RRID:AB_2532952 |
| Rabbit IgG isotype control | Vector Laboratories | Cat# I-1000;<br>RRID:AB_2336355 |
| Goat anti-mouse Alexa Fluor 568 | Invitrogen | A11004;<br>RRID:AB_2534072 |
| Goat anti-rabbit Alexa Fluor 568 | Invitrogen | A11011;<br>RRID:AB_143157 |
| Goat anti-rabbit Alexa Fluor 488 | Invitrogen | A11008;<br>RRID:AB_143165 |
| **Chemicals, peptides, and recombinant proteins** |  |  |
| PBS (without calcium and magnesium) | Corning | 21-040-CV |
| HBSS (with calcium, magnesium, and without phenol red) | Corning | 21-023-CV |
| StemPro™ Accutase™ Cell Dissociation Reagent | Gibco | A1110501 |
| Recovery™ Cell Culture Freezing Medium | Gibco | 12648010 |
| Anti-Adherence Rinsing Solution | STEMCELL Technologies | 07010 |
| Collagenase from *Clostridium histolyticum* | Sigma-Aldrich | C5138 |
| DNase I, bovine pancreas | Millipore | 260913 |
| Bovine pituitary extract | Corning | CLS354123 |
| Cholera toxin | Sigma Aldrich | C8052 |
| Retinoic acid | Sigma Aldrich | R2625 |
| Bovine serum albumin, fraction V, heat shock treated | Fisher BioReagents | BP1600-100 |
| Insulin from bovine pancreas | Sigma Aldrich | I6634 |
| Holo-transferrin bovine | Sigma Aldrich | T1283 |
| Epidermal growth factor | Fujifilm Biosciences | 100-26 |
| HyClone™ penicillin-streptomycin | Cytiva | SV30010 |
| Amphotericin B | Sigma Aldrich | A2942 |
| Avantor Seradigm Premium Grade Fetal Bovine Serum | Avantor | 97068-085 |
| DMEM/F12 (without phenol red and with 4 mM L-glutamine) | Gibco | 21041025 |
| Trypan blue solution, 0.4% (w/v) in PBS | Corning | 25-900-CI |
| Hoechst 33342 | Invitrogen | H1399 |
| Syto Deep Red Nucleic Acid Stain | Invitrogen | S34900 |
| VECTASHIELD Vibrance® Antifade Mounting Medium | Vector Laboratories | H-1700 |
| Goat serum | Sigma Aldrich | G9023 |
| 4% paraformaldehyde in PBS | Thermo Scientific Chemicals | J61899.AK |
| Triton X-100 | Sigma Aldrich | T-8532 |
| Dimethyl sulfoxide | Fisher BioReagents | BP231-100 |
| **Other** |  |  |
| AggreWell™400 24-well plate | STEMCELL Technologies | 34411 |
| Costar® 24-well Cell Culture Plate (TC-treated) | Corning | 3524 |
| Cell lifter | Corning | 3008 |
| Dissecting tissue forceps | Fisherbrand | 13-812-41 |
| Nunc™ 15 mL Conical Centrifuge Tubes (sterile, polypropylene) | Thermo Scientific | 339650 |
| 50 mL Easy Reader™ Plastic Centrifuge Tubes | Fisherbrand | 06-443-21 |
| 40 µm Sterile cell strainer | Fisherbrand | 22-363-547 |
| Petri dishes | United Scientific | 470340-170 |
| 35 mm Glass Bottom Dish with 10 mm Micro-well | Cellvis | D35-10-0-N |
| Mr. Frosty™ Freezing Container | Thermo Scientific | 5100-0001 |
| Centrifuge | Eppendorf | 5810R |
| Forma Series 3 Water Jacketed CO<sup>2</sup> Incubator | Thermo Scientific | 4110 |
| Zeiss LSM 900 confocal microscope | ZEISS | N/A |

## Materials and equipment

**DMEM/F12:** DMEM/Ham’s F12 (without phenol red and with 4 mM L-glutamine, and 15 mM HEPES), 1% penicillin and streptomycin, and 1 μg/mL amphotericin B.

**DMEM/F12 supplemented with 10% FBS:** DMEM/F12 and 10% Fetal Bovine Serum

**Note:** Can be prepared in advance and stored for up to 1 month at 4°C.

**Collagenase solution:** Dissolve 1 mg of Collagenase from *Clostridium histolyticum* in 1 mL of DMEM/F12. Add DNase I to a final concentration of 500 units/mL.

**Note:** Collagenase solution should be used on the same day and kept on ice.

### Cell Dissociation Medium

| Reagent | Final concentration | Amount |
| --- | --- | --- |
| PBS (without calcium and magnesium) | 50% | 12.3 mL |
| StemPro™ Accutase™ Cell Dissociation Reagent | 50% | 12.3 mL |
| DNase I (2 MU/mL) | 2,500 units/mL | 31.25 µL |
| **Total** | **n/a** | **25 mL** |

**Note:** Cell Dissociation Medium should be used on the same day and brought to room temperature prior to use.

### Incubation Medium

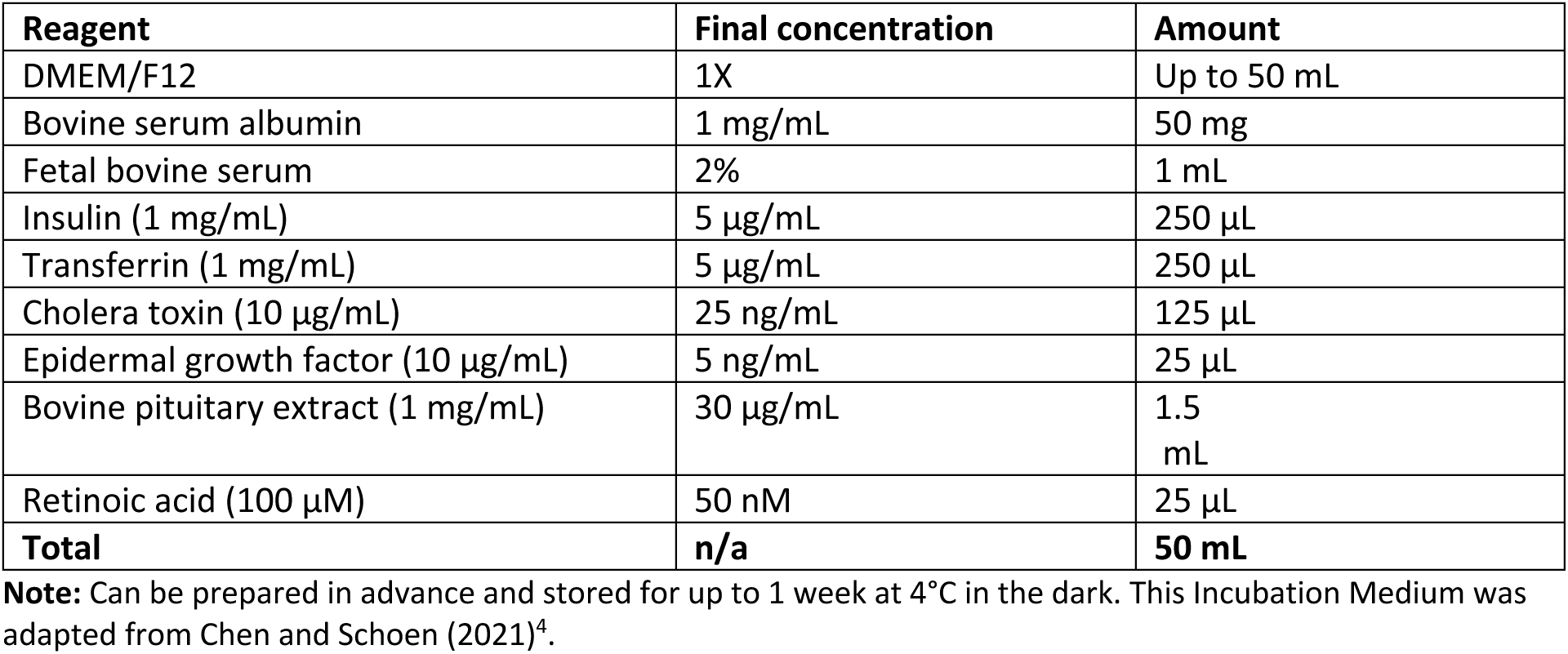

| Reagent | Final concentration | Amount |
| --- | --- | --- |
| DMEM/F12 | 1X | Up to 50 mL |
| Bovine serum albumin | 1 mg/mL | 50 mg |
| Fetal bovine serum | 2% | 1 mL |
| Insulin (1 mg/mL) | 5 µg/mL | 250 µL |
| Transferrin (1 mg/mL) | 5 µg/mL | 250 µL |
| Cholera toxin (10 µg/mL) | 25 ng/mL | 125 µL |
| Epidermal growth factor (10 µg/mL) | 5 ng/mL | 25 µL |
| Bovine pituitary extract (1 mg/mL) | 30 µg/mL | 1.5 mL |
| Retinoic acid (100 µM) | 50 nM | 25 µL |
| **Total** | **n/a** | **50 mL** |

**Note:** Can be prepared in advance and stored for up to 1 week at 4°C in the dark. This Incubation Medium was adapted from Chen and Schoen (2021)<sup>4</sup>.

**CRITICAL:** Retinoic acid is a teratogen and requires specific handling precautions. Prepare stock solutions in DMSO or 100% ethanol in a biosafety cabinet wearing appropriate PPE. Store stock solutions protected from light at -20°C.

**CRITICAL:** Cholera toxin is a potent bacterial toxin that can cause severe biological effects upon inhalation, ingestion, or exposure to the skin/mucosa. For more information read the appropriate safety data sheet. Handle with appropriate personal protective equipment inside a biosafety cabinet.

## Step-by-step method details

### Harvesting oviduct epithelial cells

#### Timing: ∼2 h

This section describes the dissection of the porcine female reproductive tract and the isolation of oviduct epithelial cells after collagenase pretreatment to preserve cell viability.

**Note:** We recommend harvesting cells from up to 6 tracts simultaneously with a single technician. Certain steps are labor-intensive, and prolonged handling may reduce cell viability.

1. Place the reproductive tract from a non-cycling prepubertal gilt on bench paper and orient it with the connective tissue facing downward (**Figure 1A**).

2. Locate the oviduct and carefully trim away surrounding connective tissue using surgical scissors (**Figure 1B**).

**Note**: If connective tissue remains, it can cause the oviduct to curl and make the outer surface of the tube uneven, both of which can interfere with the subsequent scraping process.

3. Isolate the isthmus region of the oviduct (∼5–8 cm in length; **Figure 1C**).

**Note:** The isthmus can be distinguished from the ampulla by its thicker smooth muscle layer.

4. Immediately transfer both isthmuses (left and right) to a 15 mL conical tube containing PBS supplemented with 1% penicillin/streptomycin. Incubate for 5 min at room temperature to remove residual blood.

5. In a biosafety cabinet, transfer the isthmuses onto a sterile paper towel and pat dry. Spray the outside of the oviduct with 70% ethanol and allow to air dry for a few seconds to decontaminate the outer surface.

**Note:** From this point onward, perform all steps inside a biosafety cabinet and use sterile technique.

6. Using a P200 pipette, flush the lumen of the isthmus with ∼50 to 100 μL of DMEM/F12 by inserting the pipette tip into the end closest to the ampulla (**Figure 1D**).

**Note:** This will aid in removing blood, immune cells, and oviductal fluid from the lumen before collagenase treatment.

7. Transfer the oviducts to a new conical tube containing HBSS supplemented with 1% penicillin/streptomycin and incubate again at room temperature for 5 min.

8. Using a P200 pipette, fill the lumen of each oviduct with ∼50–100 μL of collagenase solution from the end closest to the ampulla.

9. Transfer the oviducts to a 6-well plate, add ∼2 mL of DMEM/F12 per well, and incubate at 37°C in a humidified incubator for 1 h.

10. During incubation, prepare 15 mL conical tubes with 2 mL of warm DMEM/F12 supplemented with 10% FBS.

11. Following incubation, transfer the oviducts to a new 6-well plate containing 2 mL/well of DMEM/F12 supplemented with 10% FBS.

12. Transfer one isthmus into a medium-sized petri dish with ∼3 mL of DMEM/F12.

13. Using dissecting tissue forceps, grasp the oviduct at the end closest to the uterotubal junction (UTJ).

14. While holding the oviduct firmly, gently and steadily apply pressure with a cell lifter along the length of the isthmus to expel epithelial cells from the lumen (**Figure 1E**).

a. Repeat this motion once or twice to release the luminal epithelial cells.

**Note:** Applying pressure starting from the ampulla and moving toward the UTJ may reduce cell yield. **Note:** Avoid excessive force when scraping the oviduct. Collagenase treatment softens the tissue and increases the risk of rupture.

**Note:** Do not over-harvest. After the initial release of epithelial cells, one or two additional passes are typically sufficient. Attempting to over-harvest may result in contamination with blood, stromal, or muscle cells.

**Note:** Use a sterile glass slide to release epithelial cells from the oviduct if a cell lifter is not available.

15. Using a P1000 pipette, carefully collect the released cell sheets and transfer them into the prepared 15 mL conical tube containing DMEM/F12 supplemented with 10% FBS.

**Note:** Collect only the cells released from the lumen of the oviduct to minimize contamination with other cells that could detach during the scraping process.

16. Repeat steps 12–15 for the isthmus from the opposite side of the tract.

17. Repeat the procedure for all remaining samples.

**Note:** Do not combine cells originating from different animals, as residual immune cells may be present in cell collection which could result in allogeneic immune responses.

**Note:** Successful isolation of oviduct cells produces large epithelial sheets in suspension with actively beating cilia.

### Isolation and dissociation of oviduct epithelial cells into small clusters and single cells

#### Timing: ∼3 h

This section details the dissociation of oviduct epithelial cell sheets into small clusters and single cells using an enzymatic treatment and two filtration steps to generate porcine oviduct epithelial cells (POECs) suitable for spheroid formation.

18. Pass the cell suspension through a 40 μm sterile cell strainer placed over a waste collection tube and gently rinse with ∼5 mL of DMEM/F12.

**Note:** The large epithelial cell sheets are retained on the top of the strainer, while isolated single cells and debris pass through into the waste tube. Discard the flow-through. Ensure the retained cell sheets on the strainer remain wet to prevent drying.

19. Invert the strainer over a fresh labeled 50 mL conical tube and flush the retained cell sheets into it using 10 mL of DMEM/F12.

20. Centrifuge at 100 × g for 8 min.

21. Resuspend the cell sheets in 2 mL of DMEM/F12 supplemented with 10% FBS and incubate at 37°C for 30 min to allow post-isolation recovery.

22. Centrifuge the epithelial cell sheets at 100 × g for 5 min.

23. Remove the supernatant, carefully resuspend the cell pellet in 5 mL of PBS, and centrifuge again at 100 × g for 8 min.

24. Remove the supernatant and resuspend the cell pellet in 5 mL of Cell Dissociation Medium.

**Note:** Cell Dissociation Medium should be equilibrated to room temperature prior to use.

25. Incubate the cells for 10 min at room temperature. Orient the tubes horizontally to improve exposure to Cell Dissociation Medium and reduce clumping.

26. Quench the dissociation reaction by adding 5 mL of DMEM/F12 supplemented with 10% FBS and 500 μL of undiluted FBS.

27. Gently mix the cells by pipetting up and down for ∼30 sec.

28. Pass the cell suspension through a new 40 μm sterile cell strainer into a new 50 mL conical tube and rinse with 5 mL of DMEM/F12.

29. Centrifuge at 100 × g for 8 min and resuspend the cell pellet in 2 mL of DMEM/F12 supplemented with 10% FBS.

30. Centrifuge at 250 × g for 8 min and resuspend the cell pellet in 2 mL of DMEM/F12 supplemented with 10% FBS.

31. Using a hemocytometer, estimate the viability and concentration of the harvested POECs.

a. Use Trypan blue solution in a 1:10 dilution to assess cell viability.

b. Evaluate the presence of ciliated cells and ciliary motility.

**Note:** Successful isolation of POECs suitable for spheroids is indicated by the presence of small epithelial cell clusters and single cells. A high proportion of ciliated cells with actively beating cilia should be observed.

**Optional:** If spheroids do not need to be generated immediately, cryopreserve the cells at this step. Resuspend the final dissociated cell pellet in Recovery Cell Culture Freezing Medium at a density of 1.7 × 10^6^ cells/mL. Transfer 1 mL aliquots into cryovials. Place the vials into a Mr. Frosty freezing container and store at −80°C for 24 h to ensure a standard cooling rate of −1°C/minute. Following this, immediately transfer the cryovials to a liquid nitrogen vapor phase tank for long-term storage.

**CRITICAL:** Cryopreserve only cells with 80% viability or greater. Lower viability rates will result in poor cell aggregation during the spheroid formation step. Additionally, cell cryopreservation decreases the proportion of ciliated cells and reduces overall cell viability. It is ideal to utilize freshly harvested POECs for spheroid formation.

### Formation of POEC Spheroids Using AggreWell plates

#### Timing: ∼48 h

This section describes the seeding of POECs into AggreWell 400 plates to generate spheroids. POECs should consist of a mixture of small cell clusters and single cells with high viability and robust cilia activity.

**Note:** Keep the cells from step 31 at 37°C while preparing AggreWell plates.

32. Add 500 μL of Anti-Adherence Rinsing Solution to each well in AggreWell 400 24-well plate.

33. Centrifuge the plate at 1,300 × g for 5 min.

**Note:** Inspect the well for air bubbles. If bubbles are present repeat step 33.

34. Remove the Anti-Adherence Rinsing Solution and add 2 mL of PBS to each well.

35. Remove the PBS and add 1 mL of DMEM/F12.

36. Remove the DMEM/F12 and add 1 mL of DMEM/F12 supplemented with 10% FBS.

37. Add 1 mL of POEC suspension (1.7 × 10^6^ cells/mL) in DMEM/F12 supplemented with 10% FBS to each well from the AggreWell plate.

**Note:** If loss or de-differentiation of cilia is a concern during assembly, standard DMEM/F12 supplemented with 10% FBS may be replaced with Incubation Medium strictly for the spheroid formation and harvesting stages (steps 36–43). This specialized medium, adapted from established epithelial differentiation protocols^4^, is utilized here as an alternative formulation to support ciliary preservation across the 48 h aggregation window. It should not be used for downstream functional assays, such as sperm-binding experiments (step 56).

**Note:** Although cryopreserved cells can be used, they typically show reduced viability and cilia activity, which may reduce spheroid quality. Freshly isolated cells are recommended.

38. Centrifuge the plate at 100 × g for 3 min.

39. Examine the plate under an inverted microscope.

**Note:** Cells should be evenly distributed within microwells and at the appropriate concentration (**Figure 2**). Cilia movement should be observable (≥ 400 × magnification).

**Figure 2.**
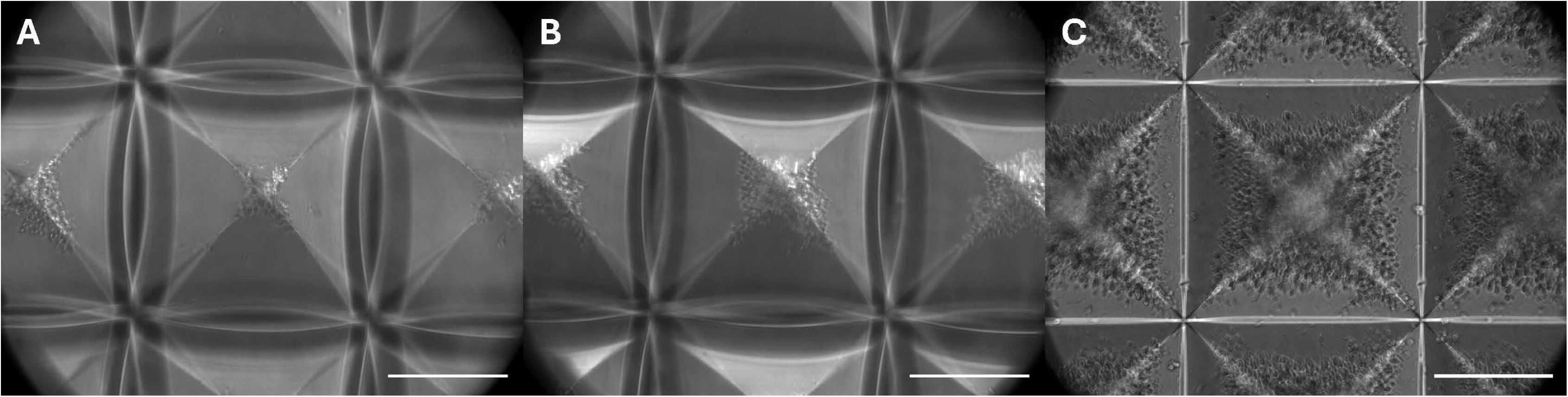
Seeding porcine oviduct epithelial cells into AggreWell 400 plates. Representative brightfield images showing cell distribution in AggreWell 400 microwells following seeding and centrifugation at different cell concentrations: (A) low cell concentration (<1.2 × 10^6^ cells/well), (B) optimal cell concentration (∼1.7 × 10^6^ cells/well), and (C) high cell concentration (>2.2 × 10^6^ cells/well). Scale bars = 200 μm.

**Note:** Handle the AggreWell plate carefully. Vigorous shaking may cause the cells to become resuspended and result in uneven settling. This may lead to inconsistent final spheroid sizes.

40. Incubate the AggreWell plate at 37°C with 5% CO_2_ for 24 h.

41. Carefully aspirate the culture media from the edge of the well without disturbing the aggregates.

42. Gradually add 1 mL of fresh DMEM/F12 supplemented with 10% FBS along the wall of each well without disturbing the aggregates.

43. Incubate for an additional 24 h to allow the spheroids to form at 37°C with 5% CO_2_ to allow for spheroid maturation.

44. Harvest the spheroids using a wide-bore P1000 pipette tip.

a. Gently aspirate the media hovering directly above the microwells to release the spheroids, then transfer the suspension to a 15 mL conical tube.

b. Add 1 mL of fresh DMEM/F12 per well to replenish the AggreWells and repeat the rinsing processes until all spheroids are collected into the tube.

c. Verify complete spheroid harvest using an inverted microscope.

45. Centrifuge the spheroids at 50 × g for 3 min. Remove the supernatant and carefully resuspend the spheroids in 2 mL DMEM/F12 supplemented with 10% FBS for each well from an AggreWell Plate harvested.

**Note:** Successful spheroid formation produces uniform spheroids with consistent size and morphology (**Figure 3**). Ciliated cells should be present, with actively beating cilia oriented outward (**Figure 4A; Methods video S1**).

**Figure 3.**
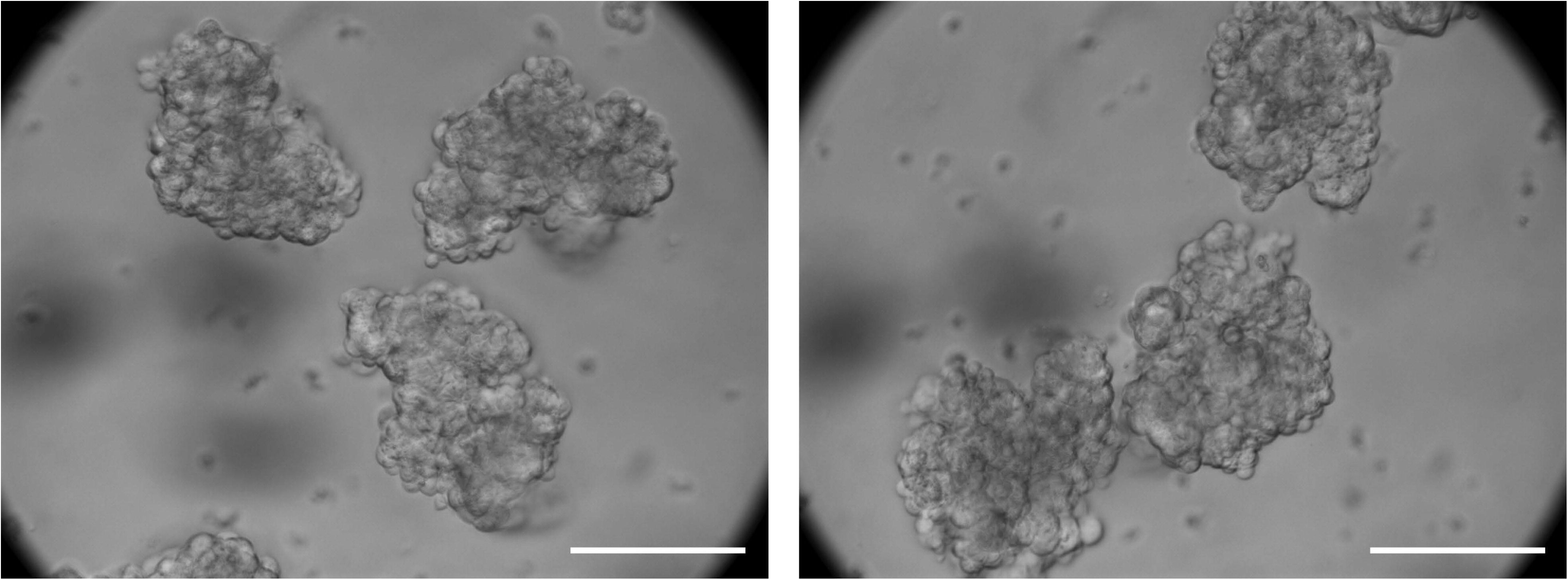
Formation of porcine oviduct epithelial cell spheroids. Representative brightfield images of porcine oviduct epithelial cell spheroids harvested after 48 h of incubation at 37°C and 5% CO2. Spheroids exhibit uniform size and morphology following aggregation in AggreWell 400 plates. Scale bar = 100 μm.

**Figure 4.**
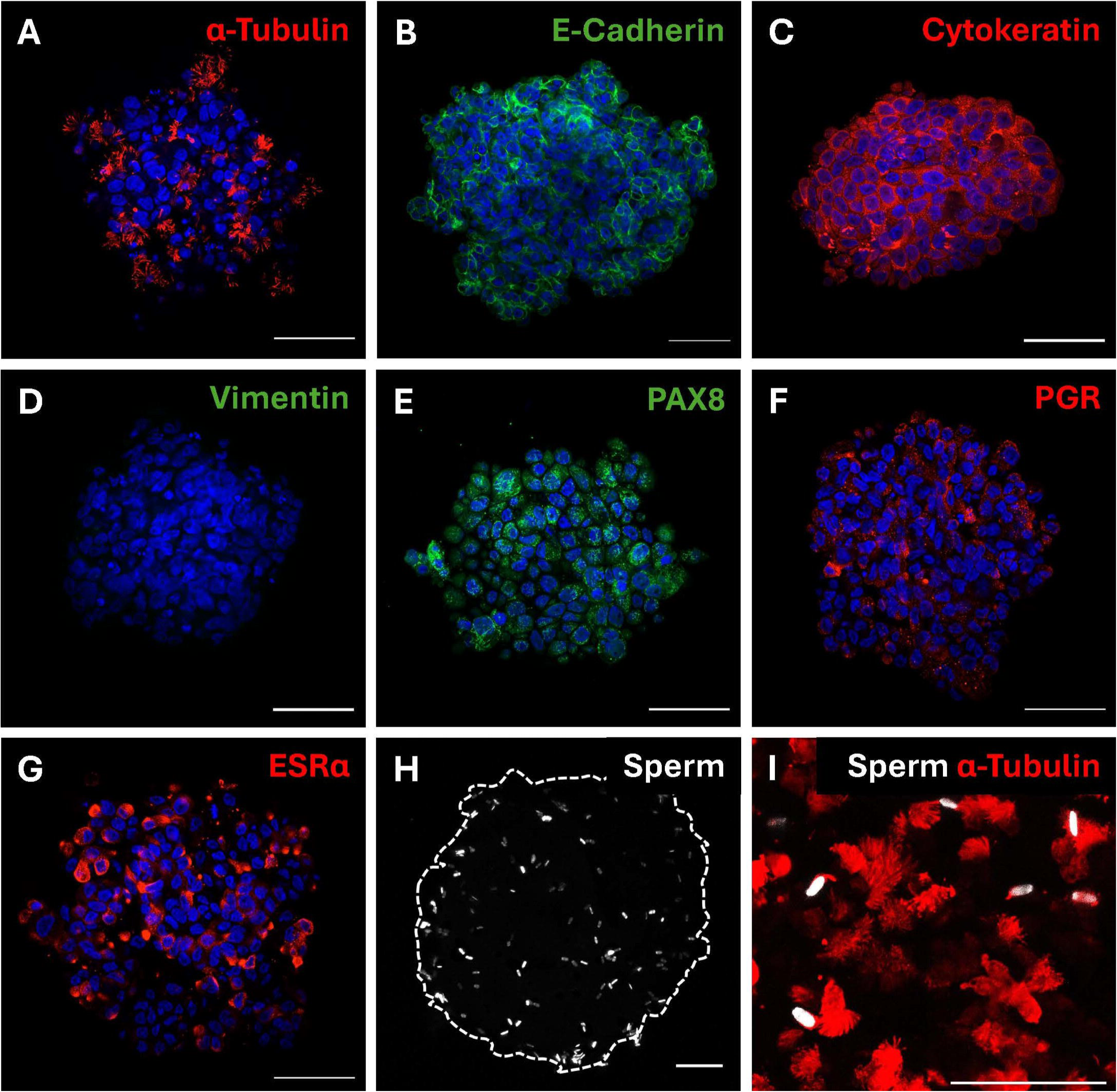
Characterization of porcine oviduct epithelial cell spheroids and sperm binding. Representative confocal images of porcine oviduct epithelial cell spheroids stained with Hoechst 33342 (blue) and immunolabeled for (A) acetylated α-tubulin (red), (B) E-cadherin (green), (C) pan-cytokeratin (red), (D) vimentin (green), (E) PAX8 (green), (F) progesterone receptor (red), and (G) estrogen receptor α (red). (H) SYTO Deep Red-labeled sperm (white) adhered to POEC spheroids following co-incubation. (I) High-magnification image showing sperm (white) bound to ciliated cells labeled with acetylated α-tubulin (red). Scale bars = 50 μm.

**Note**: The concentration of spheroids may be adjusted depending on the downstream application. **CRITICAL:** Cryopreservation of intact spheroids for later use in downstream experiments is not recommended, as freezing compromises their 3D structural integrity and cell viability. Spheroids must be generated fresh and used immediately upon harvesting.

**Note:** For downstream quantitative PCR (qPCR) analysis or RNA-seq, use at least half of the spheroids from one AggreWell to obtain sufficient RNA yield.

**Note**: For RNA extraction, always sonicate samples, even when using a strong lysis reagent such as TRIzol. Spheroids are more difficult to lyse than single cells in suspension and omitting sonication may affect the final RNA yield.

### Sperm incubation with POEC spheroids

#### Timing: ∼2 h

This section describes the incubation of POEC spheroids with sperm for sperm-binding assays and provides a practical workflow for evaluating the suitability of spheroids in sperm co-incubation experiments.

46. Add 250 μL of Anti-Adherence Rinsing Solution to each well of a 24-well plate and incubate for 5 min at room temperature. This prevents spheroid attachment to the well surface.

47. Decant the solution and rinse each well three times using 1× PBS.

48. Rinse each well once with DMEM/F12.

49. Add 900 μL of DMEM/F12 supplemented with 10% FBS to each well.

50. Add 1 mL of POEC spheroid suspension (approximately 500 spheroids/mL) to each well.

**Note:** For sperm-binding assays, fewer spheroids may be sufficient.

51. Fluorescently label sperm by adding 1 μL of SYTO Deep Red (2,000x stock) to 2 mL of extended semen. Incubate at 37°C for 30 min.

52. Wash the sperm by diluting the 2 mL of stained semen in 10 mL of DMEM/F12 and centrifuging at 600 × g for 8 min.

53. Remove the supernatant, resuspend the sperm pellet in 5 mL of DMEM/F12, and centrifuge at 600 × g for 5 min.

54. Resuspend the sperm pellet in 1 mL of DMEM/F12 supplemented with 10% FBS.

55. Adjust the sperm concentration to ∼30 × 10^6^ sperm/mL.

56. Add 100 μL of sperm suspension (approx. 3 × 10^6^ sperm) to each well containing POEC spheroids.

57. Incubate sperm with the POEC spheroids for 1 h at 37°C.

**Note**: Use a 1 h incubation to obtain more consistent results, although sperm binding typically peaks at approximately 30 min.

**Note:** Extended sperm incubation or excessive shaking may cause spheroids to adhere to one another, which can complicate downstream imaging and analysis.

58. Following incubation, gently transfer spheroids to a new well containing DMEM/F12 supplemented with 10% FBS. Incubate for an additional 2 min to remove loosely bound sperm.

**Note:** When handling the spheroids, use a P20 pipette with a wide-bore tip to minimize mechanical stress.

59. Transfer the spheroids to a second fresh well containing DMEM/F12 supplemented with 10% FBS for an additional 2 min wash.

60. Transfer the spheroids to 4% paraformaldehyde and incubate at 37°C for 30 min.

61. Transfer the spheroids to a Hoechst 33342 solution (10 μg/mL in PBS) and incubate for 10 min at room temperature.

62. Wash spheroids twice by sequentially transferring them to two wells containing 1x PBS for 5 min each.

63. Add 4 μL of VECTASHIELD Mounting Media to the center of a 35 mm glass-bottom dish.

**Note:** Using non-setting mounting media or using too much mounting media will prevent the spheroids from being immobilized while imaging. This could cause drifting of the spheroids during image acquisition.

64. Carefully transfer the spheroids to the center of the mounting medium.

**Note:** While a P20 pipette with a wide-bore tip should be used for general handling of spheroids to minimize the carryover residual buffer, switching to a P10 pipette during the mounting step is recommended. Excessive PBS carryover could compromise proper curing and spheroid immobilization.

65. Incubate the glass-bottom petri dish flat at room temperature in the dark for 2 h to allow the spheroids to settle to the bottom and for mounting media to start to harden prior to imaging.

**Note:** Imaging immediately after mounting could result in spheroids not settling properly to the bottom of the dish, making it impossible to image with high-magnification oil-immersion objectives. **Note:** Avoid disturbing or moving the dishes excessively during this curing period; disruptions can cause spheroids to migrate toward the edges of the mounting media droplet, making them difficult or impossible to image.

**Note**: Following initial curing, the petri dishes can be stored flat at 4°C for imaging within 1–2 days.

66. Image the spheroids using an inverted confocal microscope (e.g., Zeiss LSM 900).

**Note:** Spheroids settle by gravity directly onto the coverslip surface of the glass-bottom petri dish. Place the dish flat onto the stage in its standard upright orientation; do not invert the dish itself. The optical configuration of an inverted microscope allows the objective lens to scan the immobilized samples from underneath through the glass bottom.

### Immunofluorescence staining of sperm-incubated POEC spheroids

#### Timing: ∼24 h

This section describes the fixation and immunostaining of POEC spheroids for fluorescence imaging. Begin this section after harvesting POEC spheroids in step 45, or after incubation with sperm in step 58.

**Note:** Additional handling of spheroids with bound sperm will inevitably result in sperm detaching throughout the series of immunostaining steps. To mitigate this loss, perform all spheroid handling using a P20 pipette equipped with a wide-bore pipette tip.

67. Transfer the harvested spheroids to a 24-well plate containing warm DMEM/F12.

68. Transfer the spheroids to a fresh well containing warm PBS and incubate for 3 min.

69. Transfer the spheroids to a fresh well containing warm 4% paraformaldehyde and incubate at 37°C for 30 min.

**CRITICAL**: Fixation must be performed at 37°C using pre-warmed solutions. Room-temperature fixation can induce cold shock, causing the 3D cell aggregates to prematurely disaggregate.

Furthermore, incubation at 37°C accelerates the penetration rate of PFA into the dense structural core of the spheroids, ensuring uniform and complete cross-linking.

70. Wash the spheroids twice by sequentially transferring them into wells containing PBS for 5 min each.

71. Transfer the spheroids to blocking buffer (5% goat serum with 0.1% Triton X-100 in PBS) and incubate for 1 h at room temperature.

72. Transfer the spheroids to a 96-well plate containing 200 μL of primary antibody diluted in blocking buffer (typically 1:200; optimize as needed) and incubate overnight at 4°C.

73. Wash the spheroids by transferring through three wells of PBS for 5 min each.

74. Transfer the spheroids to wells with secondary antibody diluted in blocking buffer (typically 1:1,000; optimize as needed) and incubate for 1 h at room temperature.

75. Transfer spheroids to Hoechst 33342 solution (10 μg/mL in PBS) and incubate for 10 min at room temperature.

76. Wash the spheroids three times in PBS for 5 min each.

77. Carefully mount the spheroids as described in steps 63–65.

78. Image the spheroids using an inverted confocal microscope (e.g., Zeiss LSM 900).

**Note:** Spheroids may also be mounted in glass-bottom 96-well plates to prevent the spheroids from being flattened. Mounting on glass slides is possible if spheroid flattening is not a concern.

## Expected outcomes

Successful implementation of this protocol should produce uniformly sized POEC spheroids within 48 h after seeding into AggreWell plates. Spheroids should exhibit outward-facing ciliated cells with actively beating cilia (**Methods video S1**). Cell viability should remain high throughout spheroid formation, and spheroids should retain structural integrity during handling and washing steps required for downstream applications – such as sperm co-incubation, immunofluorescence staining, and harvesting for qPCR. For downstream qPCR or RNA-seq analysis, intact structural morphology is expected up until the point of sample collection, at which stage complete cellular lysis and sonication are required to maximize RNA yield from the dense 3D spheroid core.

Immunofluorescence characterization of POEC spheroids demonstrates the preservation of multiple epithelial and lineage-associated markers, including E-cadherin, pan-cytokeratin, PAX8 (secretory cell marker), acetylated α-tubulin (ciliated cell marker) and the minimal expression of fibroblast marker vimentin (**Figure 4A–E**). Additionally, the POEC spheroids retain the progesterone and estrogen receptors (**Figure 4F–G**) supporting their potential responsiveness to steroid hormones. Following sperm co-incubation, abundant sperm binding to POEC spheroids is consistently observed (**Figure 4H**), particularly along ciliated regions (**Figure 4I**), supporting the functional relevance of the model for studying sperm-oviduct interactions.

To further characterize transcriptomic changes associated with spheroid formation, we performed standard 3ʹ RNA-seq (Plasmidsaurus). Bulk RNA sequencing analysis showed that the majority of isolated POECs were epithelial cells, with high expression of essential epithelial and oviduct-associated marker genes and only minimal detection of monocyte/macrophage, fibroblast, smooth muscle, and other non-epithelial lineage marker transcripts (**Figure 5A**). Consistent with active spheroid self-organization, spheroids showed elevated expression of proliferation-associated epithelial genes, such as MKI67 and MCM2, suggesting active cell proliferation during spheroid self-organization. The top differentially expressed genes in spheroids versus freshly scraped cells revealed broad transcriptional changes associated with transition from freshly isolated epithelium to 48 h *in vitro* spheroid culture, including altered expression of stress-response, epithelial remodeling, secretory, and culture-adaptation-associated genes (**Figure 5B**).

**Figure 5.**
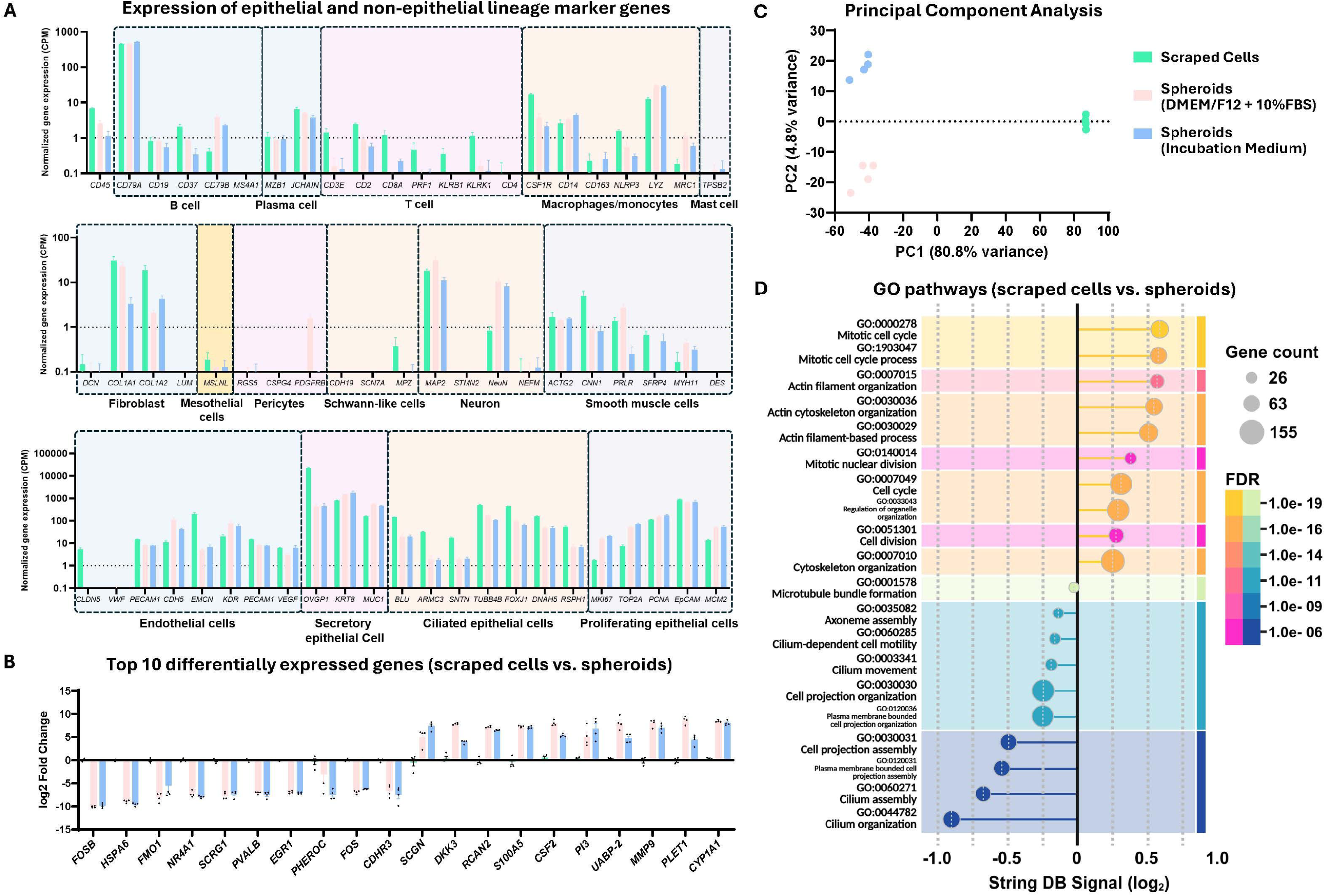
Bulk RNA sequencing analysis of freshly isolated cells and POEC spheroids. Bulk RNA sequencing was performed on freshly isolated oviduct epithelial cells (green), POEC spheroids cultured in DMEM/F12 supplemented with 10% FBS (pink), and POEC spheroids cultured in Incubation Medium (blue). (A) Cell-type marker gene expression across freshly scraped cells and POEC spheroids cultured under different conditions. The dotted line indicates 1 count per million (CPM), below which expression is generally considered background noise. (B) Top 10 down- and up-regulated genes in spheroid samples vs. scraped cells. Fold change was normalized to cells. For panels A and B, data are shown as mean ± SEM; n = 4 biological replicates. (C) Principal component analysis (PCA) showing transcriptomic clustering of the three experimental groups. (D) Gene Ontology (GO) biological process enrichment analysis using STRING-DB for differentially expressed genes between scraped cells and FBS-cultured POEC spheroids. Warm and cool colors indicate up- and downregulated genes after culture, respectively. STRING-DB signal score represents a composite enrichment score that balances the observed/expected gene ratio and FDR.

Principal component analysis showed strong clustering among biological replicates within each group, supporting the reproducibility of this spheroid model (**Figure 5C**). The first principal component, which accounted for 80.8% of the total variance, separated freshly scraped cells from spheroids. Both spheroid preparations clustered together along this axis regardless of culture medium, indicating that the transition from freshly isolated epithelium to spheroid culture is the dominant transcriptional shift in the dataset. Differences between the two media conditions were comparatively minor, resolving only along PC2 (4.8% of variance) (**Figure 5C**).

Gene Ontology (GO) biological process enrichment analysis of differentially expressed genes identified pathways related to cell-cycle activity, cytoskeletal organization, and cilium organization/motility, consistent with rapid epithelial remodeling during spheroid formation and retention of ciliated epithelial features (**Figure 5D–E**). Overall, these findings indicate that POEC spheroids rapidly self-organize into a biologically active, three-dimensional, polarized epithelial model that undergoes expected culture-associated remodeling while retaining key oviduct-associated characteristics.

## Limitations

This protocol describes the generation of porcine oviduct epithelial cell spheroids using primary oviduct epithelial cells isolated from pre-pubertal gilts. Because the model relies on primary tissue rather than immortalized cell lines, biological variability among animals might affect cell yield, viability, and spheroid formation efficiency. Although RNA sequencing analysis demonstrated strong correlations between biological replicates (**Figure 5C**), variation in donor tissue quality remains an inherent limitation of primary cell-based systems.

The success of this protocol is highly dependent on the physiological state of the donor animal. While consistent spheroid formation is achieved using tissues collected from pre-pubertal gilts, isolation of epithelial cells from post-pubertal cycling animals is less reliable and often results in lower cell quality and reduced spheroid formation efficiency. Parity and stage of estrous cycle may therefore significantly influence reproducibility. Although spheroid formation can still be achieved using tissue from post-pubertal animals, efficiency and consistency are typically reduced compared with pre-pubertal donors.

The protocol is optimized for rapid spheroid formation (48 h) to preserve ciliated epithelial cells and maintain ciliary activity. Extended culture periods may result in progressive loss of ciliary activity or epithelial de-differentiation, which may affect the physiological relevance of the model for sperm-oviduct interaction studies. Media composition is also a critical factor, as changes in supplementation can alter epithelial differentiation state and transcriptomic profiles. Bulk RNA sequencing demonstrated distinct transcriptional differences between spheroids cultured in standard DMEM/F12 supplemented with 10% FBS and spheroids cultured in Incubation Medium, indicating that the phenotype of spheroids can be modulated through culture conditions. We believe this tunability provides a controllable system for studying oviduct epithelial biology and differential cellular responses to environmental cues.

Finally, transcriptomic analysis also revealed substantial differences between freshly isolated oviduct epithelial cells and spheroids cultured for 48 h. These differences likely reflect rapid adaptation to *in vitro* culture and active self-organization during spheroid formation, including changes associated with epithelial remodeling, cell-cycle activity, cytoskeletal organization, and cilia-associated programs. Thus, although POEC spheroids retain key oviduct epithelial characteristics, they should not be interpreted as fully recapitulating the immediate transcriptional state of freshly isolated oviduct tissue. This distinction should be considered when interpreting gene expression-based outcomes.

## Troubleshooting

### Problem 1

Low cell viability or reduced ciliary activity during dissociation (see steps 24–26).

Prolonged exposure to Cell Dissociation Medium reduces cell viability and impairs ciliary activity.

### Potential solution

- Reduced viability often originates from poor starting material. Ensure that harvested cells exhibit high viability prior to dissociation.
- Limit incubation time with Cell Dissociation Medium to the minimum required for effective dissociation. Overexposure can lead to excessive cell death and extracellular DNA release, which may result in the formation of viscous, gel-like aggregates and reduced viability.

### Problem 2

Cells fail to settle into AggreWell microwells after centrifugation (see steps 37–39).

After centrifugation, viable cells and small clusters should settle evenly into the microwells. Cells that remain suspended are often non-viable and may fail to incorporate into spheroids.

### Potential solution

Repeat centrifugation, step 38, to promote cell settling. If cells continue to float or fail to fully settle into the microwells, this typically indicates poor cell viability and may compromise spheroid formation. If sufficient cells are present within the microwells (see Figure 2), proceed with incubation. Floating debris and dead cells can be carefully removed during medium exchange (step 41–42).

### Problem 3

Formation of small or poorly developed spheroids (see steps 37–45).

Spheroids that are smaller than expected may result from insufficient seeding density or reduced cell viability.

### Potential solution

- Increase the initial seeding density of POECs in AggreWell plates.
- Use Figure 2 as a reference for expected microwell occupancy following seeding and centrifugation.
- Confirm that cells are evenly distributed within microwells following centrifugation.
- Avoid cryopreserved cells when possible, as cryopreservation can reduce overall viability and number of ciliated cells. Use freshly isolated POECs for optimal spheroid formation.

### Problem 4

Loss of cilia or reduced ciliary activity during culture (see steps 40–44).

Ciliated cells may progressively lose ciliary structure or exhibit reduced ciliary motility during culture.

### Potential solution

- Ensure POECs exhibit robust ciliary activity prior to spheroid formation.
- Use freshly isolated cells whenever possible.
- Avoid excessive disturbance during medium changes, as this may damage cilia or disrupt spheroid integrity.
- If longer culture periods are required, we recommend using the Incubation Medium to better support maintenance of the ciliated phenotype.

### Problem 5

Spheroids are lost due to sticking to the inside of the pipette tip during washing and staining steps (see steps 58–64 and 67–77).

Spheroids are naturally highly adherent and sample loss can occur due to sticking to the internal surface of the plastic pipette tip during spheroid handling steps.

### Potential solution

- Pre-coat the pipette tips before handling the spheroids. Pipet Anti-Adherence Rinsing Solution up and down several times in a sterile microcentrifuge tube to coat the interior surface of the tip. Rinse the tip by pipetting PBS up and down before transferring spheroids.

**Note:** Anti-Adherence Rinsing Solution may exhibit cellular toxicity. Consequently, this tip-coating technique should strictly be limited to handling spheroids that are already fixed or are intended for immediate downstream fixation.

### Problem 6

Spheroids drift during imaging (see steps 63–65 and 77–78).

Spheroids may drift during imaging or Z-stack acquisition if the mounting medium has not sufficiently stabilized.

### Potential solution

- Allow mounting medium to partially cure before imaging to reduce spheroid movement.
- Ensure that spheroids are mounted in the center of the mounting medium droplet during mounting.
- Keep mounted samples flat, undisturbed, and protected from light during the settling period to promote stabilization prior to imaging.
- Avoid excessive movement when transporting samples to the microscope for imaging.

### Problem 7

Low RNA yield from POEC spheroids for qPCR or RNA sequencing (see steps 44–45).

RNA yield from harvested spheroids may be insufficient for downstream applications such as qPCR or RNA sequencing, particularly when spheroid number or cell viability is low.

### Potential solution

- Increase the number of POEC spheroids collected per experimental group to improve total RNA yield.
- Sonicate samples after resuspending in TRIzol or the desired lysis buffer to ensure efficient lysis of spheroids.
- Ensure high cell viability prior to spheroid seeding, as low viability will reduce RNA yield and quality.
- Minimize delays between spheroid harvesting and RNA extraction to reduce RNA degradation.

## Resource availability

### Lead contact

Further information and requests for resources and reagents should be directed to and will be fulfilled by the lead contact, Dr. David J. Miller.

### Technical contact

Technical questions on executing this protocol should be directed to and will be answered by the technical contact, Leonardo M. Molina and Hao-Chun Fan.

### Material availability

This study did not generate new unique reagents.

### Data and code availability

Bulk RNA sequencing and primary data processing were performed by Plasmidsaurus (San Francisco, CA) using their standard 3ʹ RNA-seq workflow. Sequencing quality control, read alignment, transcript annotation, and gene-level quantification were conducted by the vendor. The vendor-generated gene count matrix was used for downstream analysis. The bulk RNA-sequencing data generated during this study are available in the NCBI Gene Expression Omnibus (GEO) under accession number GSE335056.

## Acknowledgments

We thank Rantoul Foods for providing porcine female reproductive tracts from gilts. Research reported in this publication was supported by the National Institutes of Health Eunice Kennedy Shriver National Institute of Child Health and Human Development under award numbers R21HD111954 to DM and AA, and NIH F31HD108959 to LM. The content is solely the responsibility of the authors and does not necessarily represent the official views of the National Institutes of Health.

## Author contributions

L.M.M. and H.C.F. designed and performed the experiments. L.M.M., H.C.F., A.M.A. and D.J.M. interpreted the results. L.M.M. wrote the first version of the manuscript, and L.M.M., H.C.F., A.M.A. and D.J.M. reviewed and edited the manuscript. All authors have read and approved the manuscript.

## Declaration of generative AI and AI-assisted technologies in the writing process

During the preparation of this work, the authors used ChatGPT and Grammarly to improve language and readability without altering the core content. After using this tool, the authors reviewed and edited the content as needed and take full responsibility for the content of the publication.

**Methods video S1. Actively beating cilia on the surface of a POEC spheroid.**

Real-time brightfield/live-cell imaging showing the active ciliary beating of differentiated epithelial cells on a freshly harvested spheroid imaged at 630X.

## References

1. Molina, L.M., Pepi, L.E., Shajahan, A., Doungkamchan, K., Azadi, P., McKim, D.B., and Miller, D.J. (2023). Siglecs in the Porcine Oviduct and Sialylated Ligands on Sperm: Roles in the Formation of the Sperm Reservoir. bioRxiv, 10.1101/2023.03.26.534240.

2. Ferraz, M.A.M.M., Henning, H.H.W., Stout, T.A.E., Vos, P.L.A.M., and Gadella, B.M. (2017). Designing 3-Dimensional In Vitro Oviduct Culture Systems to Study Mammalian Fertilization and Embryo Production. Ann Biomed Eng 45, 1731–1744. 10.1007/s10439-016-1760-x.

3. Ferraz, M.A.M.M., Rho, H.S., Hemerich, D., Henning, H.H.W., van Tol, H.T.A., Hölker, M., Besenfelder, U., Mokry, M., Vos, P.L.A.M., Stout, T.A.E., et al. (2018). An oviduct-on-a-chip provides an enhanced in vitro environment for zygote genome reprogramming. Nat Commun 9, 4934. 10.1038/s41467-018-07119-8.

4. Chen, S., and Schoen, J. (2021). Using the Air–Liquid Interface Approach to Foster Apical–Basal Polarization of Mammalian Female Reproductive Tract Epithelia In Vitro. In Next Generation Culture Platforms for Reliable In Vitro Models : Methods and Protocols Methods in Molecular Biology., T. A. L. Brevini, A. Fazeli, and K. Turksen, eds. (Springer US), pp. 251–262. 10.1007/978-1-0716-1246-0_18.

5. Ford, M.J., Harwalkar, K., and Yamanaka, Y. (2022). Protocol to generate mouse oviduct epithelial organoids for viral transduction and whole-mount 3D imaging. STAR Protocols 3, 101164. 10.1016/j.xpro.2022.101164.

6. Menjivar, N.G., Gad, A., Thompson, R.E., Meyers, M.A., Hollinshead, F.K., and Tesfaye, D. (2023). Bovine oviductal organoids: a multi-omics approach to capture the cellular and extracellular molecular response of the oviduct to heat stress. BMC Genomics 24, 646. 10.1186/s12864-023-09746-y.

7. Kessler, M., Hoffmann, K., Brinkmann, V., Thieck, O., Jackisch, S., Toelle, B., Berger, H., Mollenkopf, H.-J., Mangler, M., Sehouli, J., et al. (2015). The Notch and Wnt pathways regulate stemness and differentiation in human fallopian tube organoids. Nat Commun 6, 8989. 10.1038/ncomms9989.

8. Lõhmussaar, K., Kopper, O., Korving, J., Begthel, H., Vreuls, C.P.H., van Es, J.H., and Clevers, H. (2020). Assessing the origin of high-grade serous ovarian cancer using CRISPR-modification of mouse organoids. Nat Commun 11, 2660. 10.1038/s41467-020-16432-0.

9. Schmaltz, L., Prudhomme, T., Tsikis, G., Reynaud, K., Mérour, I., Mermillod, P., and Saint-Dizier, M. (2024). Sperm binding to oviduct epithelial spheroids varies among males and ejaculates but not among females in pigs. Theriogenology 219, 116–125. 10.1016/j.theriogenology.2024.02.022.

10. Pranomphon, T., Mahé, C., Demattei, M.-V., Papillier, P., Vitorino Carvalho, A., Reynaud, K., Almiñana, C., Bauersachs, S., Parnpai, R., Mermillod, P., et al. (2024). Characterization of oviduct epithelial spheroids for the study of embryo–maternal communication in cattle. Theriogenology 217, 113–126. 10.1016/j.theriogenology.2024.01.022.

